# A robust method for RNA extraction and purification from a single adult mouse tendon

**DOI:** 10.1101/247379

**Authors:** M Grinstein, HL Dingwall, RR Shah, TD Capellini, JL Galloway

**Author notes:** co-first authorship (M Grinstein, HL Dingwall).

## Abstract

**Background:** Mechanistic understanding of tendon molecular and cellular biology is crucial towards furthering our abilities to design new therapies for tendon and ligament injuries and disease. Recent transcriptomic and epigenomic studies in the field have harnessed the power of mouse genetics to reveal new insights into tendon biology. However, many mouse studies pool tendon tissues or use amplification methods to perform RNA analysis, which can significantly increase the experimental costs and limit the ability to detect changes in expression of low copy transcripts.

**Methods:** Single Achilles tendons were harvested from uninjured, contralateral injured, and wild type mice between 3-5 months of age, and RNA was extracted. RNA Integrity Number (RIN) and concentration were determined, and RT-qPCR gene expression analysis was performed.

**Results:** After testing several RNA extraction approaches on single adult mouse Achilles tendons, we developed a protocol that was successful at obtaining high RIN and sufficient concentrations suitable for RNA analysis. We found that the RNA quality was sensitive to the time between tendon harvest and homogenization, and the RNA quality and concentration was dependent on the duration of homogenization. Using this method, we demonstrate that analysis of *Scx* gene expression in single mouse tendons reduces the biological variation caused by pooling tendons from multiple mice. We also show successful use of this approach to analyze *Sox9* and *Col1a2* gene expression changes in injured compared with uninjured control tendons.

**Discussion:** Our work presents a robust, cost-effective, and straightforward method to extract high quality RNA from a single adult mouse Achilles tendon at sufficient amounts for RNA-seq and RT-qPCR. We show this can reduce biological variation and decrease the overall costs associated with experiments. This approach can also be applied to other skeletal tissues as well as precious human samples.

## Introduction

Tendon injuries are common problems for active individuals and the aging population (Kaux J-F 2011). Treatment options include physical therapy and surgical intervention, but pain and limited mobility often persist, making complete restoration of tendon function challenging (Nourissat et al. 2015). Our current understanding of the molecular and cellular pathways regulating tendons during homeostasis, healing, and aging are limited. Several studies using large animal models such as sheep, rabbits, and rats have provided important information about tendon injury, biomechanics, surgical techniques, and bioengineering strategies for tendon repair (Voleti 2012). Other studies have used mouse genetics to gain an understanding of the molecular and cellular response of tendons to acute injuries, changing load environments, and in gene loss-of-function models (Mendias et al. 2008), (Dunkman et al. 2014; Dyment et al. 2014), (Howell et al. 2017), (Wang et al. 2017). The mouse system offers unique advantages for implementing mechanistic studies of tendon biology as they permit genetic lineage tracing and conditional knockout strategies, and they can be housed simply and in large numbers to improve sample sizes for functional studies. Even with inbred mouse strains, inter-animal variation can affect the conclusions drawn from gene expression analyses (Sultan et al. 2007), (Watkins-Chow & Pavan 2008). Therefore, of the use of several biological replicates of tendon tissues obtained from individual mice for RNA analysis is essential for furthering our mechanistic understanding of tendon biology.

Mature tendons are comprised of type I collagen, which are arranged in a highly ordered hierarchical manner along the long axis of the tissue (Kannus 2000). Tendon cells lie between these organized fibrils and are surrounded by a hydrophilic, glycoprotein-rich ground substance (Kannus 2000) (Yoon & Halper 2005), (Bi et al. 2007). This dense, fibrous, water-rich matrix that surrounds the tendon cells poses a significant challenge for the acquisition of high-quality RNA. In addition, tendons have low cell density compared with other tissues such as muscle or liver, resulting in minimal RNA yield per gram of tissue (Kannus 2000; Reno et al. 1997).

Previous studies have described protocols for RNA extraction from human or larger mammalian animal models such as rabbit (Ireland & Ott 2000), (Reno et al. 1997), but analyzing RNA from small animal models such as mouse can be more difficult. This has led to several different strategies for achieving RNA yield and quality sufficient for gene expression analysis by RT-qPCR or RNA-seq. RNA amplification methods have permitted gene expression analysis of single injured and uninjured tendons (Dunkman et al. 2014), but this can be prohibitively expensive for analyzing a large number of samples or target genes, currently possible using the mouse system. In addition, studies in other tissues have shown that amplification can lead to biased results and increased false negative rates, especially for low- and medium-copy transcripts (Dunkman et al. 2014). Mendias and colleagues, (Mendias et al. 2008; Mendias et al. 2012) and Nielson and colleagues (Nielsen et al. 2014) have performed expression analysis on a single mouse Achilles or plantaris tendon in different loss-of-function mouse models or in altered loading conditions. However, this approach is not widespread in the literature and the studies, although reporting a good 260/280 ratio, do not report on the RNA integrity as they mainly performed RT-qPCR. However, there are examples of many studies that pool a large number of tendons (e.g., 12-20 individual tendons) (Bell et al. 2013), (Trella et al. 2017). Not only does this increase the mouse cohort size and experimental costs, but it can also enlarge the inter-individual variation, which may explain some of the large variability in transcript abundance that was found in subsets of their gene expression analysis (Trella et al. 2017). Lastly, other studies have focused on tendon-derived cell populations such as tendon stem/progenitor cells (Bi et al. 2007). This approach will result in robust RNA yields, but it queries a cell population that has been expanded in culture and could have altered transcriptomic and epigenomic states compared with that of the native tendon tissue.

The various technical limitations associated with obtaining high-quality, high-yield RNA enlarges the cohorts of mice needed for statistical analysis, and hinders the use of RT-qPCR or functional genomic assays such as RNA-seq on single adult mouse tendons. Here, we present a robust, low-cost, and straightforward RNA isolation protocol that enables the isolation of high-integrity RNA from a single mouse Achilles tendon. We show that pooling tendon samples inflates biological variance estimates for gene expression data in RT-qPCR analysis. We apply this method to analyze injured and contralateral uninjured tendons and demonstrate the detection of significant and reproducible gene expression changes. In addition, this method can be used to purify high quality RNA from other musculoskeletal tissues, making it easily adaptable to multiple connective and skeletal tissue types, or from difficult to obtain tissues from humans or other organisms.

## Methods

### Mouse Studies

Achilles tendons were collected from wildtype C57BL/6 mice between 3-5 months of age (Jackson Laboratories 00664, n = 30 total). To compare gene expression levels between injured and uninjured Achilles tendons in the same mouse, excisional Achilles tendon injuries were performed using a 0.3 mm biopsy punch as described (Beason et al. 2012). The incision was closed with 6-0 Ethilon nylon sutures and the tendons were harvested 30 days after injury for analysis. Mice were housed, maintained, and euthanized according to American Veterinary Medical Association guidelines. All experiments were performed according to our Massachusetts General Hospital Institutional Animal Care and Use Committee (IACUC: 2013N000062) approved protocol.

### RNA Extraction and Purification

Dissected Achilles tendons were placed immediately into 1.5 ml tubes containing 500 μl of TRIzol reagent (Invitrogen Cat# 15596026) and high impact zirconium 1.5 mm beads (30-40 beads per tube, D1032-15 Benchmark). Samples were homogenized immediately in two, 180-second rounds of bead beating at 50 Hz (BeadBug microtube homogenizer). Samples were then moved directly to dry ice or -80°C for longer storage up to 6 months.

To extract RNA, the samples were thawed on ice followed by a 5 minute incubation at room temperature. Samples were quickly spun in the sample tubes and the homogenate was moved to a new Eppendorf tube, leaving behind the beads and residual tissue. Next, a chloroform extraction was performed, using double the recommended amount, which has been shown to increase RNA yields in small samples (Macedo 2014). 100 μl of chloroform was added to the homogenate and vortexed well for approximately 1 minute. The Trizol/chloroform mixture was then moved to a 1.5 ml MaXtract high density tube (Qiagen Cat No. 129046), incubated at room temperature for 2-3 minutes, and spun ≥12,000 x g at 4°C for 15 minutes. MaXtract tubes contain a sterile gel that forms a barrier between the RNA-containing aqueous phase and the Trizol/chloroform upon centrifugation at 4°C, thus minimizing carryover of organic solvents leading to an overall reduction in sample contamination. After centrifugation, the aqueous phase was transferred to a clean 1.5 ml Eppendorf tube and an equal volume of 100% ethanol was added to the aqueous phase and mixed well. At this stage, the RNA/ethanol mix was typically stored at -80°C. We have found that brief incubation of this mixture at -80°C improved the total RNA yield, yet it is not required.

RNA purification was next performed using the ZR Tissue & Insect RNA MicroPrep kit (Zymo Research R2030) or the Direct-Zol systems (Zymo Research R2050, R2060). Based on typical tendon yields, the ZymoSpin IC spin columns are optimal for use with RNA extracted from single tendons as these columns can purify up to 5 μg of RNA in as little as 6 μl eluate. However, this protocol also has been successfully used with ZymoSpin IIC columns, which require a larger elution volume. After adding the RNA/ethanol mix to the spin column, the standard Zymo purification protocol was used with the following modifications. First, a 15-minute on-column DNase I treatment was added to minimize genomic DNA contamination. An extra wash step was included to improve sample purity. Prior to elution, columns were spun for an additional 2 minutes at maximum speed to remove residual ethanol. RNA was eluted in 15 μl RNase/DNase free water that was pre-warmed to 55-60°C to maximize the RNA recovery from the spin column. RNA concentration was measured via fluorometric quantitation (Qubit HS RNA assay, Invitrogen, CAT# Q32852) and sample quality was determined by spectrophotometric analysis (NanoDrop 2000c, ThermoFisher Scientific) as well as capillary electrophoresis (2100 Bioanalyzer, Agilent). The final RNA product was stored at -80C for RT-qPCR analysis.

### RT-qPCR, Data Analysis, and Statistics

100 ng total RNA was reverse transcribed with oligo(dT)_20_ primers using the SuperScript IV First Strand Synthesis System (Thermo Fisher 18091050) and a no-reverse transcriptase control was included for every sample. A total of 2 ng cDNA template was amplified for 40 cycles in each SYBR green qPCR assay (Applied Biosystems 4367659) using a final primer concentration of 200 nM. All assays were performed in technical triplicate using either a LightCyclerII 480 (Roche; pooled samples) or a StepOnePlus Real Time PCR system (Applied Biosystems; injury samples). Three independent biological samples were run per condition for both sets of RT-qPCR. *Gapdh* was used as the reference gene for all samples (see Table 1 for primer sequences).

**Table 1.**
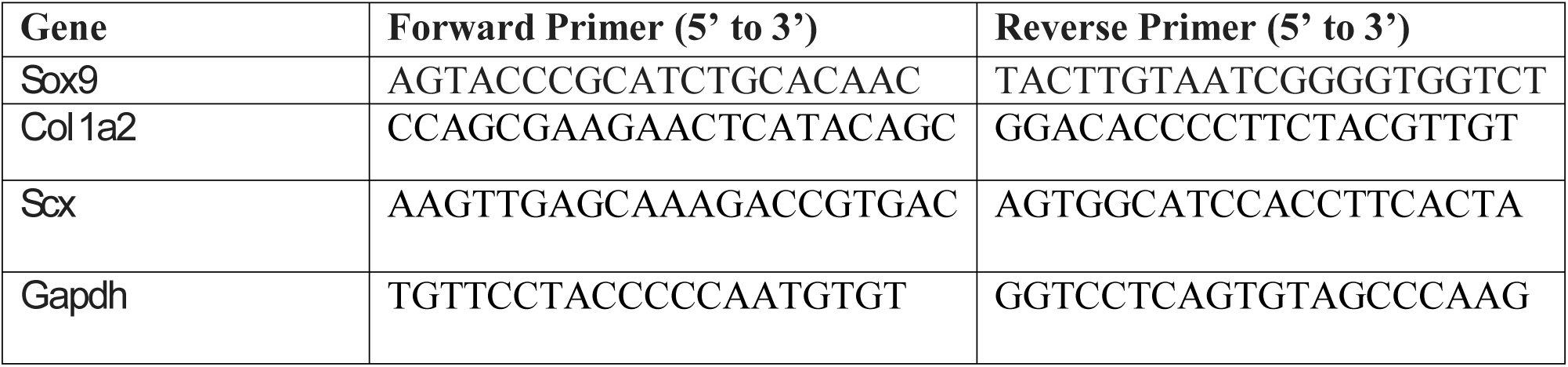
Primers used for RT-qPCR

All analyses were conducted in R (R Core Team 2017). For the pooling experiment, summary statistics were calculated for *Scleraxis* (*Scx*) and *Gapdh* technical and biological replicate cycle threshold (C_T_) values independently. Variance estimates for *Scx* ΔC_T_ relative expression were calculated using standard error propagation techniques. Relative expression values for *Collagen Ia2* (*Col1a2*) and *SRY-Box9* (*Sox9*) were calculated for the injury analysis using the ΔΔC_T_ method (Livak & Schmittgen 2001) and injury samples were normalized to their corresponding uninjured contralateral controls. Statistical differences between injured and uninjured samples from three biological replicates (n=3 mice) were analyzed via Welch’s t-test (Welch 1947) on the ΔC_T_ values.

## Results

Several tissue disruption methods were tested in order to achieve optimal RNA quality and quantity from a single mouse tendon. Among those tested were enzymatic digestion, cryogenic grinding (manual and mill), shearing with a handheld homogenizer (i.e., rotor-stator), and bead beating. Capillary electrophoresis was performed on purified RNA using a Bioanalyzer RNA Nano chip (Agilent). RNA integrity number (RIN), a quantification of degradation, was calculated by the accompanying Agilent software based on the electropherogram for a given sample; a RIN of 10 indicates completely intact RNA whereas a RIN of 1 indicates severely degraded RNA. Enzymatic digestion produced intact RNA (RIN > 7), but low RNA yield (≤ 1ng/μl). Cryogenic grinding and handheld homogenizer dissociation methods resulted in low yield (≤ 5ng/μl) and poor RNA integrity (RIN ≤ 3). Bead beater homogenization was found to produce the best results in terms of RNA quality (i.e., RIN >= 6.5) and quantity (>= 50 ng/ul), and minimized carryover between samples. Additionally, bead beating was easily combined with standard TRIzol extraction and commercially available purification methods.

To further evaluate our bead beating homogenization method, we performed additional experiments examining the level of degradation that occurs prior to homogenization as well as during homogenization. To address the former, single Achilles tendons from similarly aged mice were left on ice following dissection for up to 9 minutes before homogenization in the bead beater. The shortest time between dissection and homogenization (0-30 seconds) yielded more intact RNA (RIN = 6.5) while longer wait times resulted in more degraded RNA (9 minutes processing time RIN = 5.4; Figure 1). This demonstrates that measurable degradation can occur prior to sample homogenization, and occurs with increases in time after dissection on the order of only minutes (Figure 1). Therefore, processing the dissected tendon(s) immediately following dissection is essential for preserving RNA integrity. We next tested how the duration of bead beating affects RNA quality by varying homogenization times of single and four pooled Achilles tendons. Samples were homogenized for 30 seconds, 60 seconds, 180 seconds, or 360 seconds (in two consecutive rounds of 180 seconds; Figure 2 A, B). RNA from samples homogenized for less than 60 seconds suffered more degradation than those that underwent longer homogenization times (Figure 2B), indicating incomplete homogenization of the tissue during the shorter bead-beating periods. Homogenization times longer than 360 seconds did not improve RNA quality, and in some cases caused further degradation.

**Figure 1.**
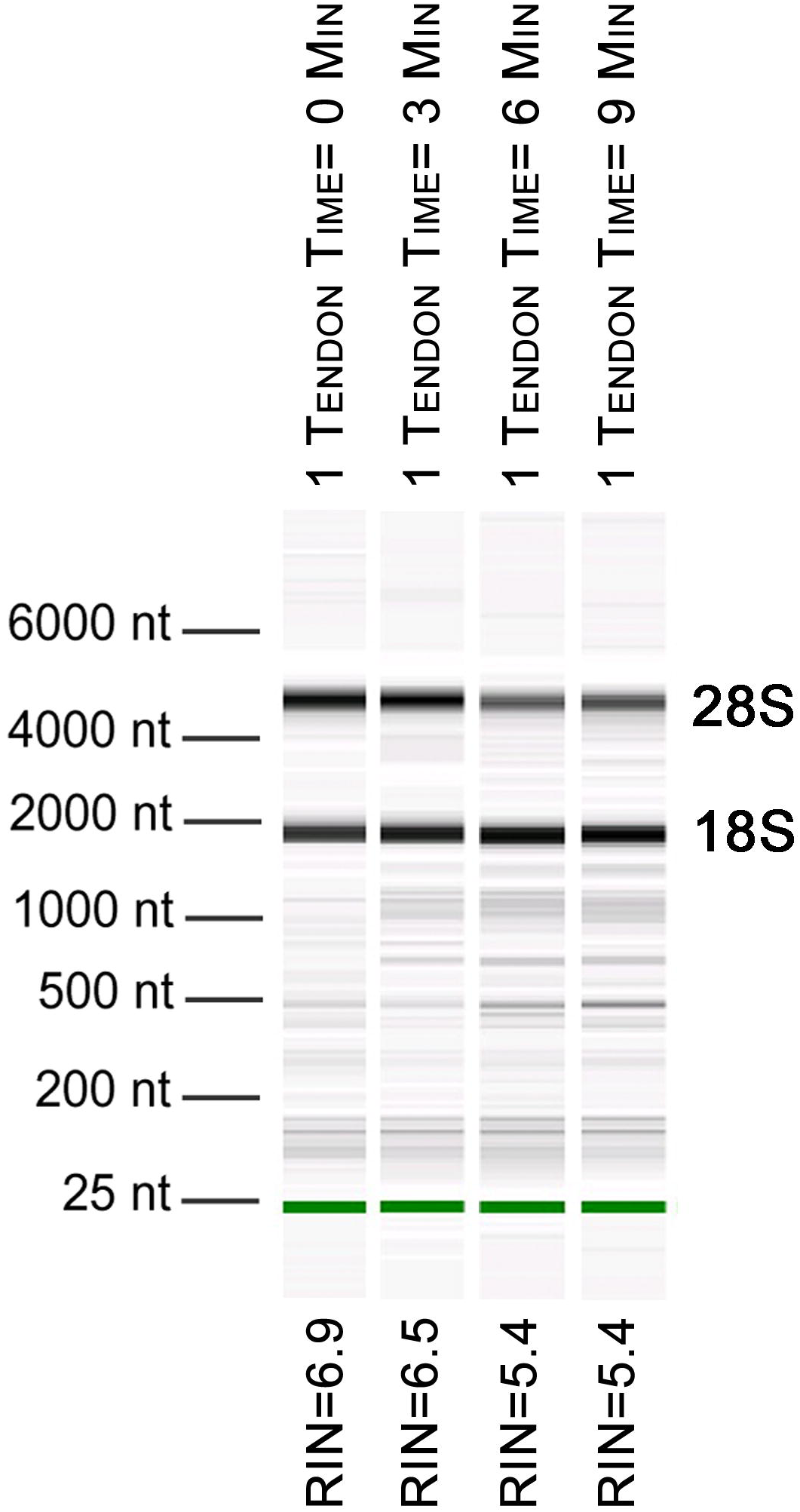
Length of time between dissection and processing affect RNA integrity. Electropherogram digital gel via Bioanalyzer shows integrity of RNA isolated from single Achilles tendons that were kept on ice for various lengths of time (0, 3, 6, 9 minutes) before homogenization. All were homogenized for 360 seconds. Longer wait times prior to homogenization reduce RNA quality. 18S and 28S are indicated and the green band is a marker.

**Figure 2.**
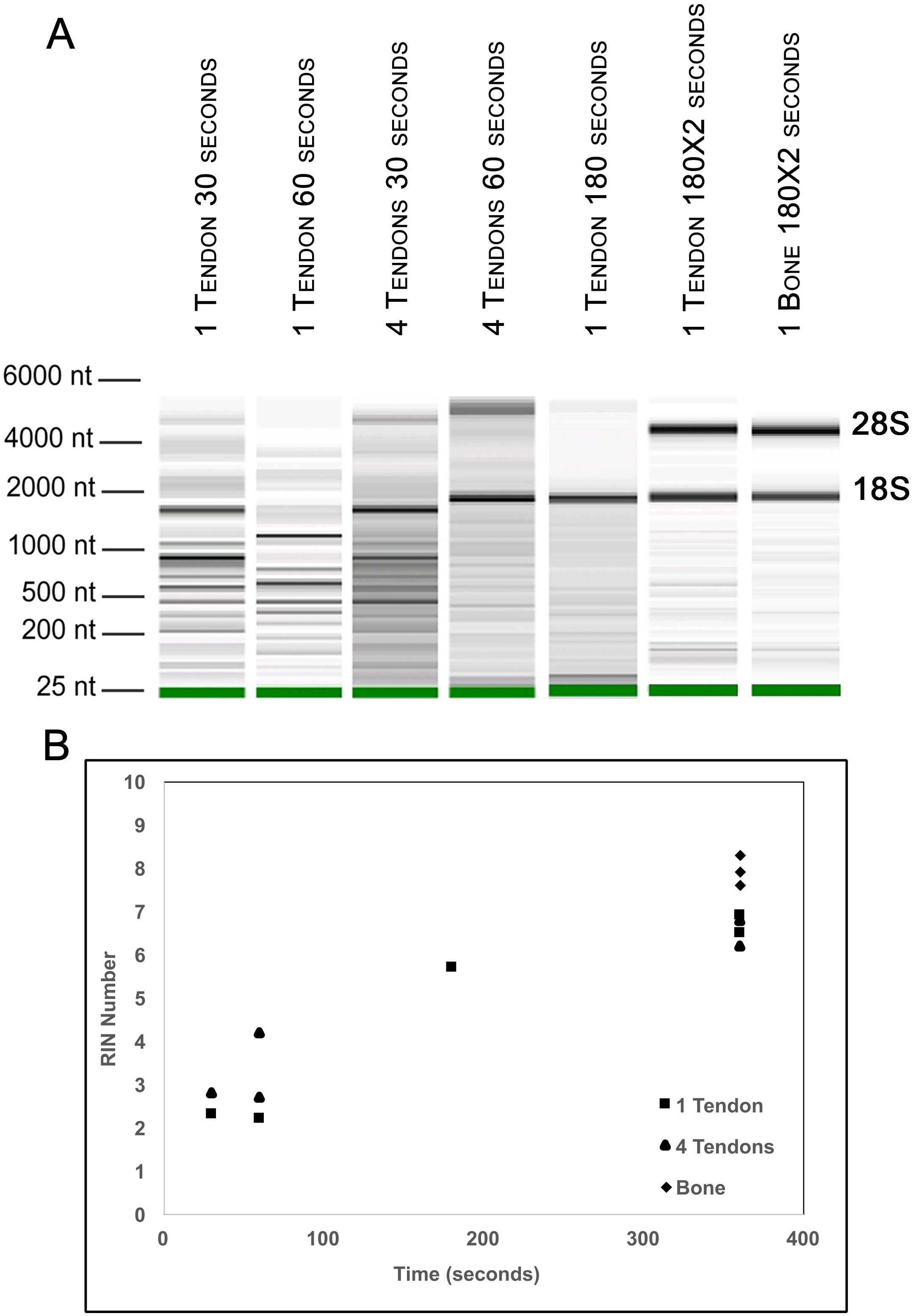
Optimization of homogenization regime. Single Achilles tendons and pools of four tendons were subjected to different durations of bead beating homogenization: 30, 60, 180, and 360 (in two rounds of 180) seconds. The electropherogram digital gel shows that the longest beating time resulted in the most intact RNA, as evidenced by the strong 28s and 18s bands with 360 seconds (A). RIN values called by Agilent software also show the improvement in quality with longer beating time (B). More than 360 seconds showed no appreciable improvement in RNA integrity (data not shown).

To test whether pooling tendons from multiple individuals into one sample prior to homogenization influences RNA integrity, we measured RNA quality from single Achilles tendons as well as pools of differing sizes (2, 4, 6, and 8 tendons, n = 3 biological replicates per pooling level; Figure 3 A, B). Electropherograms and RIN measurements show that RNA from all pooling levels suffer levels of degradation similar to single Achilles samples (Figure 3A, B). Therefore, pooling tendons from multiple individuals is not protective against RNA degradation; the only measure that improved with increased pool size was RNA yield (Figure 3C). To determine if pooling multiple samples affects gene expression measurements, we evaluated gene expression in single and differentially pooled tendon samples described above (n = 3 per pooling level) via RT-qPCR. Although we find no gain in RNA quality from pooling, treating pools of tendons from multiple individuals as single biological replicates results in larger standard deviations in C_T_ measurements in assays for *Scx* and *Gapdh* (Figure 4). This leads to larger sample variance for larger pools, driven by differences in ΔC_T_ between biological replicates within a group, which impedes the detection of small gene expression changes. Such increases in variance for pooled versus single samples have also been reported for RNA-seq datasets (Rajkumar et al. 2015).

**Figure 3.**
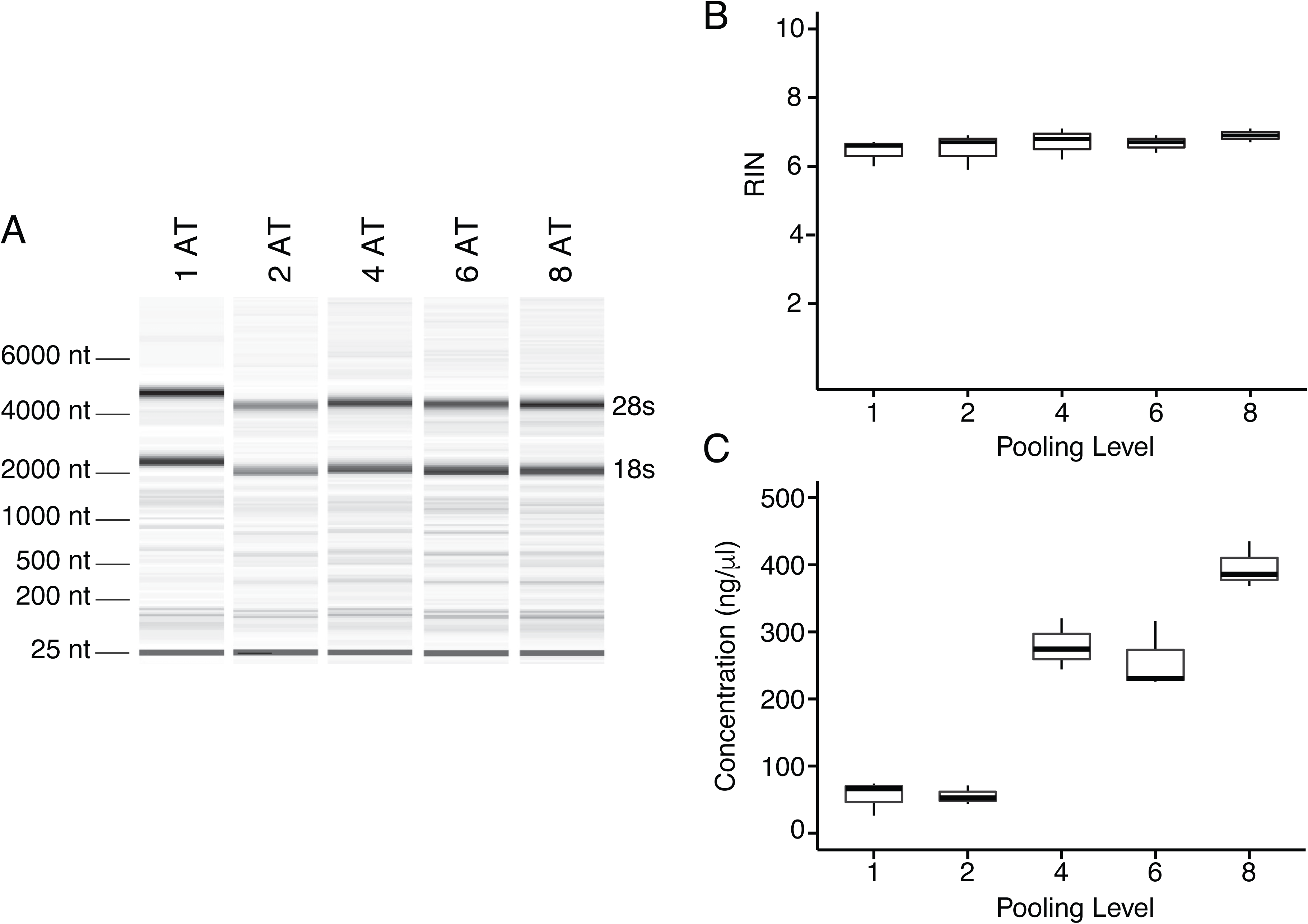
Tendon pooling affects RNA quality and yield. Representative electropherogram digital gel of RNA from different-sized pools of Achilles tendons demonstrates high integrity RNA across all samples (A). Called RINs for pools (n = 3 per pool) demonstrates that RNA quality from a single tendon is comparable to that from pools of tendons. Sample RINs are sufficiently high for use in RNA-seq gene counting and differential expression analysis for as low as one Achilles tendon (B). Concentration of RNA from single or pooled tendons increases with tendon number (n = 3 per pool) (C). The middle line represents the median, the box is quartiles 2 and 3 interquartile range (IQR), and whiskers are 1.5 x IQR (B, C).

**Figure 4.**
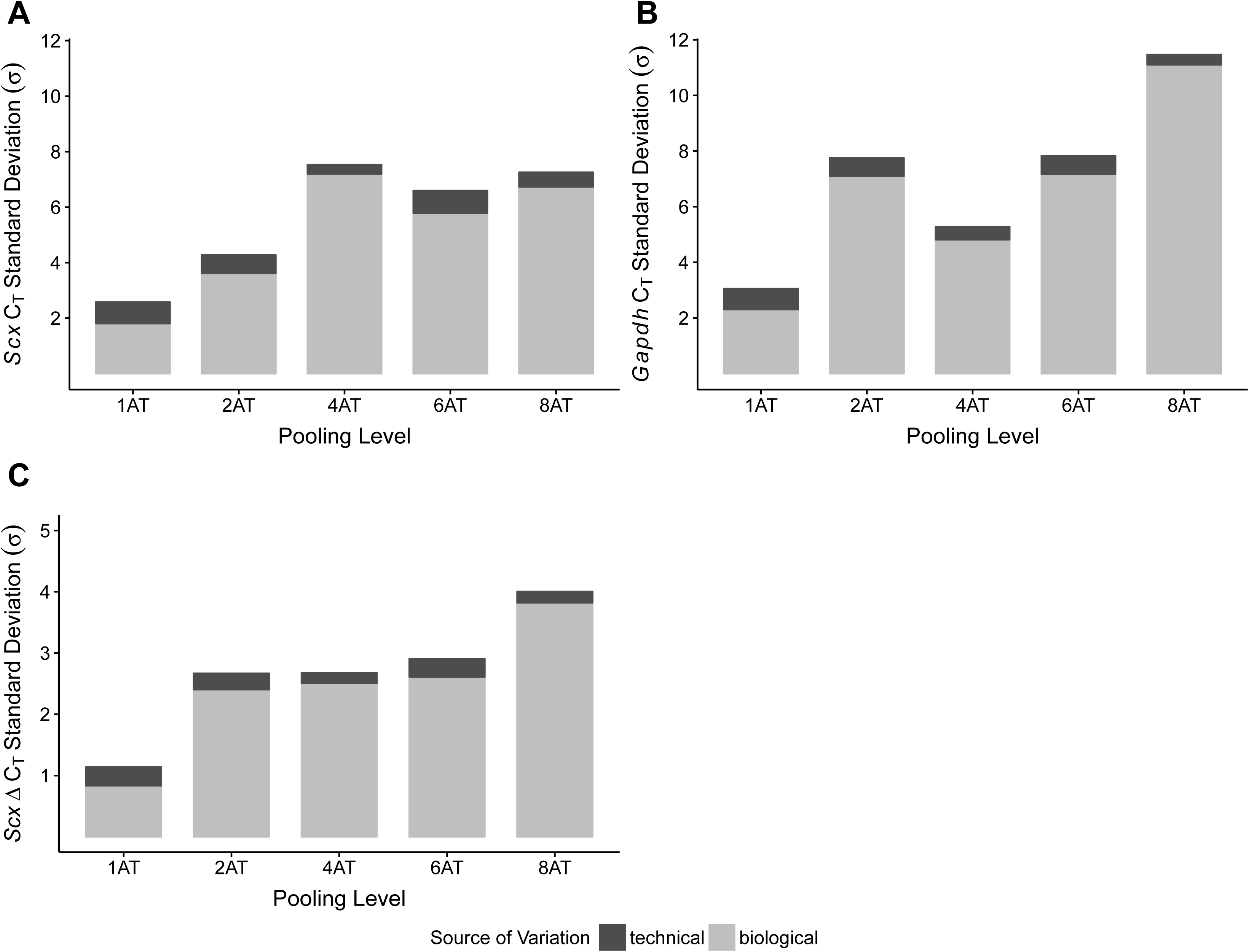
Sample pooling affects estimates of sample variance in RT-qPCR. C_T_ standard deviations for *Scx* (A) and *Gapdh* (B) measurements were calculated for the technical replicates (n = 3 repeat measurements; dark grey) and for biological replicates (n = 3 independent samples; light grey), separately. ΔC_T_ was calculated by normalizing *Scx* C_T_ values to *Gapdh.* Technical (dark grey) and biological (light grey) variance estimates were calculated separately (C). All measures (A-C) show that biological variance increases as number of individuals contributing to a pool increases.

To validate the performance of the RNA obtained using this protocol, we performed RT-qPCR for *Sox9* and *Col1a2* expression on single Achilles tendons at 30 days following an acute excision Achilles tendon injury. All samples were obtained from single injured and contralateral uninjured Achilles tendons from the same mouse. Using this protocol, we found significantly increased expression of *Sox9* and *Col1a2* in injured Achilles tendons compared with their uninjured contralateral counterparts (p < 0.05 for *Sox9* and P< 0.01 for *Col1a2*; Figure 5). These results are consistent with previous studies showing increased expression of *Sox9* and *Col1a2* following tendon injury (Guerquin et al. 2013) (Zhang & Wang 2013), and also show that our method is robust to detect gene expression changes in single tendon samples.

**Figure 5.**
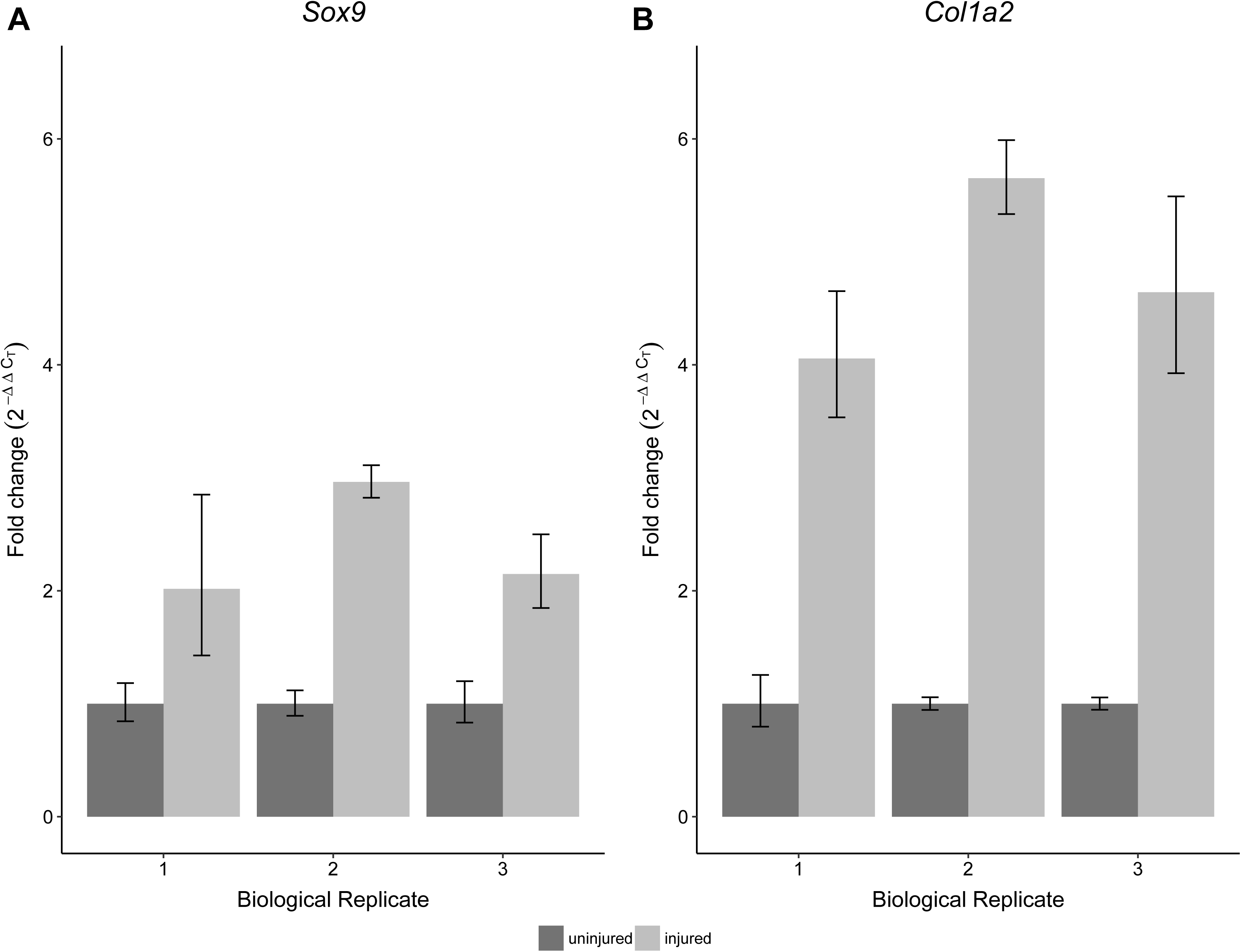
Sensitivity and reproducibility of RT-qPCR on single tendon RNA. RT-qPCR of *Sox9* (A) and *Col1a2* (B) expression of injured Achilles tendons relative to the contralateral tendon of the same mouse at 30 days post injury. A Welch’s t-test shows that both *Sox9* and *Col1a2* expression was significantly different in the injured condition compared to the control tendons (n = 3 biological replicates, (p < 0.05 for *Sox9* and P< 0.01 for *Col1a2*).

## Discussion and Conclusions

Obtaining high quality RNA from tendons can be challenging, and this can limit the direction and scope of studies focused on analysing adult mouse tendon tissues. Whereas a few studies have used single tendons without amplification, many other studies have used amplification or pooling of greater than 12 samples to detect gene expression changes. Both approaches can be expensive due to the high costs associated with amplification kits for multigene analysis or the number of mice used for one biological replicate. Dissociation, followed by culture and expansion of tendon-derived cells can yield greater RNA concentrations of high quality, but such approaches cannot be used to study gene expression changes after injury. The approach we described above provides a straightforward method to consistently obtain high yields of RNA from one Achilles tendon of sufficient quality to perform RT-qPCR analysis without amplification. In addition, the reported RIN scores are acceptable for standard RNA-seq differential expression analysis.

Our analysis also uncovered key steps that are integral towards generating high RIN and concentration from the single tendon samples. In particular, we find that the time from dissection to homogenization and storage can significantly impact the quality of the RNA, causing measurable degradation. In this regard, *even* small delays on the order of minutes could affect overall RNA quality, which could greatly affect differential gene expression analysis. In addition, the duration of homogenization is important for maximizing RNA yield and quality. Homogenization times that are too short or long can result in dramatically different RIN and concentrations regardless of the level of sample pooling.

Similar to previous RNA-seq studies, our RT-qPCR analysis of single and pooled tendon samples revealed that pooling increases the variance of gene expression measurements (Rajkumar et al. 2015). It has been argued that pooling samples from multiple individuals into single biological replicates results in biological averaging and is therefore an appropriate, and even useful, practice in gene expression studies via microarray (Kendziorski et al. 2005) However, genes, which are lowly expressed or exhibit subtle differences between conditions, would require a larger sample size of pools to achieve adequate statistical power, which would further inflate mouse and reagent cost for RT-qPCR, microarray, or RNA-seq analyses (Shih et al. 2004). This analysis also highlights the problem of performing RT-qPCR comparisons on a single pool per group (run in technical triplicate), under the assumption that the within-sample variation is representative of the biological variation among all animals of that group. Variance calculated from technical repeats does not estimate biological variance within each group, and is not an appropriate practice. The technical variation arises from noise due to measurement error and therefore is unrelated to biological variation (Kitchen et al. 2010; Vaux et al. 2012), necessitating the use of multiple pools for any statistical analysis.

Our tendon RNA extraction method is a robust protocol for obtaining high quality RNA for gene expression assays. It decreases the number of mice required for analysis and avoids extra amplification steps, making it straightforward, cost-effective, and easily accessible to researchers new to the tendon field. By providing a means for reproducibly analyzing one Achilles tendon, this method also reduces measurement error associated with pooling tendons from multiple individuals. Moreover, our protocol permits the use of internal comparisons between a limb that has undergone experimental manipulation (e.g., injury or unloading) and the contralateral control limb within the same animal. In addition to facilitating larger-scale RT-qPCR studies, we believe this method will make high dimensional gene expression analysis such as RNA-seq accessible to more researchers in the tendon and other musculoskeletal biology fields, thus opening new frontiers in tendon biology.

## Acknowledgements

We would like to thank the Harvard University Bauer Core Facility and MGH Center for Comparative Medicine for their services. J.L.G, H.L.D., M.G., and T.D.C. were supported by AR071554 NIAMS/NIH. J.L.G. and R.R.S. were supported by the American Federation of Aging Research and the Harvard Stem Cell Institute. M.G. was supported by Human Frontiers Science Program Fellowship. T.D.C was supported by the Milton Fund and Dean’s Competitive Fund (Harvard University).

